# Hypothalamic extraretinal photoreceptor Opsin3 regulates water balance, body temperature and motor activity

**DOI:** 10.1101/2022.07.28.501815

**Authors:** Soledad Bárez-López, Ghadir Elsamad, Paul Bishop, Daniel Searby, Yvonne Kershaw, Becky Conway-Campbell, David Murphy, Michael P Greenwood

## Abstract

Opsin 3 (Opn3) is a light-sensitive protein present throughout the mammalian brain, although its functions in this organ are currently unknown. Since *Opn3* encoded mRNA increases in the supraoptic nucleus (SON) of the hypothalamus in response to hyperosmotic stimuli, we have explored the role of Opn3 in this nucleus. By *in situ* hybridization, we have demonstrated that *Opn3* is expressed in neurones expressing both arginine vasopressin and oxytocin in the rat SON, suggesting that OPN3 functions in both neuronal types. Gene profiling analysis by RNA-seq suggested that neuropeptide production is impaired when knocking down *Opn3* in the rat SON, as confirmed by quantitative PCR and immunolabelling. Knocking down Opn3 in the rat SON altered physiological parameters including water intake, body temperature and motor activity. Altogether the data indicates that Opn3 in the SON is involved in the regulation of a number of neuropeptides and other proteins that participate in water homeostasis, body temperature and motor activity.

## INTRODUCTION

Synchronisation to periodic cues such as food/water availability and light/dark cycles is crucial for homeostasis. Modern lifestyles disturb our diurnal rhythms, most acutely by jetlag, but more insidiously by shift work and exposure to artificial light at night (ALAN), with consequential increased susceptibility of night-time workers to diabetes, obesity, cardiovascular disease, and depression (Richter et al., 2021). It is therefore essential to explore the mechanisms of light-regulated physiology and behaviour in order to improve health and wellbeing.

Traditionally, it was believed that opsins in the rods and cones and a subset of intrinsically photosensitive retinal ganglion cells were the only light-sensitive proteins in mammals. Following the discovery of the first extraretinal light-sensitive proteins in thornback ray and chicken pineal glands (Vigh-Teichmann et al., 1983; Okano et al., 1994), Encephalopsin or Opsin 3 (Opn3) was the first extraretinal opsin identified in the mammalian brain (Blackshaw and Snyder, 1999) suggesting that light may play a broader functional role than previously recognised.

Opn3 is a seven-transmembrane domain G-protein-coupled receptor that is activated by blue light (Koyanagi et al., 2013; Barreto Ortiz et al., 2018) and can mediate light-dependent (Barreto Ortiz et al., 2018; Sato et al., 2020) but also light-independent processes (Ozdeslik et al., 2019). Opn3 has been found to activate Gi/Go-type G proteins leading to decreases in cAMP (Koyanagi et al., 2013; Ozdeslik et al., 2019). In the embryonic mouse brain, Opn3 is observed from E9.5 in the ganglia that will develop into the trigeminal, facial, and vestibulocochlear cranial nerves and, throughout brain development, its distribution increases in several motor-sensory tracts (Davies et al., 2021). In the adult brain, *Opn3* RNA (Blackshaw and Snyder, 1999) and Opn3 protein (Nissila et al., 2012; Olinski et al., 2020) are clearly detected in the thalamus, the cerebellum, portions of the frontal cortex, the hippocampus, the anterior medial preoptic areas and several hypothalamic nucleus including the paraventricular nucleus (PVN) and the supraoptic nucleus (SON). However, more than two decades after its discovery, Opn3 remains one of the least characterised opsins and its brain functions are still unknown.

In recent transcriptome studies, we observed that *Opn3* mRNA expression in the SON is consistently modulated in response to osmotic hypo- and hypertonic challenges (Greenwood et al., 2015; Dutra et al., 2021; Pauza et al., 2021). The SON is a hypothalamic nucleus of the hypothalamo-neurohypophysial system (HNS), a neurosecretory apparatus responsible for the production of the peptide hormones arginine vasopressin (AVP) and oxytocin (OXT). The HNS consists of two distinct populations of magnocellular neurones (MCNs) in the SON and PVN that separately synthesise either AVP or OXT. MCN axons project to blood capillaries of the posterior pituitary gland neuro-vascular interface (Mecawi Ade et al, 2015; Murphy et al, 2016). To counteract hyperosmotic imbalance, such as that evoked by chronic dehydration or salt-loading (Greenwood et al., 2015a; Greenwood et al, 2014; Murphy et al., 2016), the HNS releases AVP and OXT into the bloodstream (Bargmann, 1966) to regulate water balance. AVP is also involved in cardiovascular control by mediating vasoconstriction (Rascher, 1985) and by controlling sympathetic outflow towards the cardiovascular system (Lozic et al., 2018). Moreover, MCNs from the SON also send extra-neurohypophysial collateral projections to other brain areas that can potentially regulate behavioural responses to physiological challenges (Grinevich and Ludwig, 2021) including social behaviour (Meyer-Lindenberg et al., 2011), the stress response and energy balance (Ding and Magkos, 2019).

Given the wealth of knowledge on the anatomy and physiology of this discrete hypothalamic nucleus, in this work, we have explored the role of Opn3 in the SON. We found that *Opn3* in the SON is expressed in both AVP and OXT neurones and by knocking down *Opn3* in this structure we found changes in water intake, body temperature and motor activity, suggesting a role for OPN3 in metabolic homeostasis.

## MATERIAL AND METHODS

### Animals

All experimental procedures involving animals were performed in strict accordance with the provision of the UK Animals (Scientific Procedures) Act (1986). The study was designed following the ARRIVE (Animal Research: Reporting of In Vivo Experiments) guidelines (Kilkenny et al., 2010) and carried out under Home Office UK licences (PPL 30/3278 and PPL PP9294977). All the protocols were approved by the University of Bristol Animal Welfare and Ethical Review Board.

Male Sprague-Dawley rats weighing 250–300 g (Envigo) were maintained under a 12:12 light-dark cycle, lights on at 8.00 am, (12:12 reverse cycle for animals undergoing physiological determinations, lights on at 10.00 pm) at a constant temperature of 21-22°C and a relative humidity of 40%-50% with *ad libitum* access to food and water. Animals were housed with environmental enrichment consisting of nesting material, cardboard tube, and a chew block. Rats used for RNAscope *in situ* hybridization and RNA-seq analysis were housed in groups of 2 or 3. Rats used for physiological analyses were singly housed to allow for accurate food and water intake measures. To improve animal welfare, single-housed animals were grouped together in a ball pit once a week for a minimum of 1 hour to allow for social interaction and play. The water deprivation (WD) protocol involved the removal of water for 72 hours for RNAscope analyses (PPL 30/3278) and for 48 hours (PPL PP9294977) for body temperature and motor activity measurements, with *ad libitum* access remaining for controls.

### Experimental design

4 control and 4 WD rats were used for RNAscope analysis and for immunofluorescence based on previous experience (Barez-Lopez et al., 2022). 5 control and 5 *Opn3* knockdown SON samples were used for RNA-seq analysis based on studies on power analysis and sample size estimation for RNA-Seq differential expression (Ching et al., 2014). 6 control and 7 rats where *Opn3* was knocked down in the SON (Opn3KD) were used for physiological studies based on previous findings in which knocking down another protein involved in water homeostasis in the SON (Caprin2) altered water intake when animals were salt loaded (Konopacka et al., 2015). Power calculations were conducted using the power calculation tool available through the Experimental Design Assistant (EDA) tool from the NC3Rs (Percie du Sert et al., 2017) considering a 4.46 effect size in water intake (ml/100 g b.m.) after 2 days of salt loading, a SD of 2.14, with a significance threshold (α) of 0.05 and statistical power (1-β) of 0.90, which recommended n = 6. Taking into account the complexity of the successful stereotaxic delivery of adeno-associated virus (AAV) vectors to knockdown *Opn3* in the SON, 8 animals were subjected to surgery to obtain 5 successful controls and *Opn3* knockdowns for RNA-seq, 6 to obtain 4 brains for immunofluorescence and 20 to obtain at least 12 (6 control and 6 Opn3KD) rats for physiological analysis.

Bias-reducing measures for food and water intake determination included the random distribution of control and Opn3KD rat cages using Research Randomizer (https://www.randomizer.org/) and blinding of the researcher taking the measurements until the end of the study.

### RNAscope in situ hybridization

Control euhydrated and 72-hour WD rats were euthanised by cranium striking followed by guillotine decapitation. Brains were removed and immediately frozen in powdered dry ice. Fresh-frozen brains were sliced into 16 µm coronal sections and directly mounted on Superfrost Plus slides (Thermo Fisher Scientific, Waltham, MA, USA) and stored -80 C°. RNAscope *in situ* hybridization was performed as described (Barez-Lopez et al., 2022) using the probes Rn-AVP-C2 (Advanced Cell Diagnostics, 401421-C2) targeting 20 – 525 of rat *Avp* gene, Rn_OPN3 (Advanced Cell Diagnostics, 578411) targeting 418 – 1702 of rat *Opn3* gene, and Rn-OXT-C3 (Advanced Cell Diagnostics, 479631-C3) targeting 3 – 493 of rat *Oxt* gene.

### AAV production, purification, and titration

A Opn3 short hairpin RNA (shRNA) (GGGACAGGCCAAAGAAGAAAG) and a non-targeting shRNA sequence (AATTCTCCGAACGTGTCACGT (Greenwood et al., 2020)), used as a control, were cloned into HuSH shRNA GFP AAV Cloning Vector (pGFP-A-shAAV Vector; TR30034, Origene). The efficiency of the specific *Opn3* shRNA was tested in HEK293T/17 cells (human embryonic kidney cell line, CRL-11268, ATCC) transiently overexpressing rat *Opn3*. Virus particles were produced as previously described (McClure et al., 2011). Briefly, HEK293T cells were transiently transfected with the ITR-flanked transgene together with three separate packaging plasmids including pHelper Vector (340202, Cell Biolabs), pAAV-RC1 Vector (VPK-421, Cell Biolabs), and pAAV-RC2 Vector (VPK-422, Cell Biolabs) using Lipofectamine 3000 Transfection Reagent (11668019, Invitrogen) according to the manufacture’s specifications. 72 hours after transfection, cells were harvested and lysed with 150 mM NaCl, 20 mM Tris pH 8.0, 0.5% sodium deoxycholate and 50 units per ml of benzonase nuclease. Viral particles were purified using HiTrap heparin columns (GE HealthCare Life Science), concentrated with Amicon ultra-4 centrifugal filters to 100 µl and diluted with an equal volume of sterile phosphate buffered saline (PBS). Titers of AAV vector stocks were determined by direct qPCR amplification (Aurnhammer et al., 2012).

### Viral injections

Preliminary experiments were performed to assess the efficiency of AAV-shOpn3-eGFP and AAV-shNT-eGFP transduction *in vivo*. AAV-shOpn3-eGFP and AAV-shNT-eGFP were injected into the SON as described (Konopacka et al., 2015). Briefly, 1 μl containing 1.59 × 10^9^ AAV-shOpn3-eGFP viral particles or 1 μl containing 1.31 × 10^9^ AAV-shNT-eGFP particles were injected into the SON by stereotaxic delivery at coordinates 1.3 mm caudal, ±1.95 mm lateral, and −8.8 mm ventral to bregma. For RNA-seq analysis and immunofluorescence, rats were injected with AAV-shOpn3-eGFP in one SON and with AAV-shNT-eGFP in the other SON as in (Greenwood et al., 2022). For physiological studies, rats were injected bilaterally in the SON with AAV-shOpn3-eGFP or AAV-shNT-eGFP. Successful delivery to the target region and *Opn3* knockdown efficiency were determined by *Opn3* and *Gfp* expression by quantitative PCR analysis in each individual SON. Only those samples with an *Opn3* knockdown efficiency of above 50% were used for RNA-seq analysis (average knockdown efficiency of 72.58%, ranging from 55.62% - 89.84%). Data from physiological analysis was retrospectively chosen from rats with an average of 77.21% *Opn3* knockdown efficiency.

### RNA-seq

Two weeks following unilateral administration of AAV-shOpn3-eGFP in one SON and AAV-shNT-eGFP in the other SON, rats were euthanised by cranium striking followed by guillotine decapitation. Brains were removed and immediately frozen in powdered dry ice. All sample collections were performed between 9.00 am and 12.00 pm. SON samples were collected using a 1-mm micropunch (Fine Scientific Tools, 18035-01) from 100 μm brain coronal sections in a cryostat as described (Greenwood et al., 2014). RNA from individual SONs was extracted with RNeasy Mini Kit (74106, Qiagen) according to manufacturer’s instructions and eluted in a final volume of 30 µl. RNA quality and integrity was assessed with Agilent 2100 Bioanalyzer where the average RNA integrity Numbers (RIN) value was 9.15 (ranging from 8.9 – 9.5). Poly(A)-selected mRNA was sequenced (Source Bioscience). Poly(A) enriched bulk RNA-sequencing libraries were constructed using Illumina TruSeq Stranded mRNA kits. Libraries were loaded onto lanes of an Illumina NextSeq flowcell and sequenced using 75bp paired-end (PE) runs. Each sample generated >35million PE reads. Read alignment was performed using STAR (Dobin et al., 2013) (version:020201) against the Rattus norvegicus genome (Rn6). We used FeatureCounts (v1.6.2) (Love et al., 2014) to generate read counts, using the ENSEMBL annotations for reference. We identified genes that are differentially expressed (DE) using DESeq2 (v1.32.0) in R. The analysis was sufficiently powered (n = 5) to reduce the false discovery rate, and to enable systems level analysis. Differentially expressed genes (DEGs) with p adjusted (padj) values of <0.05 are considered significant. The data underlying the transcriptomic analyses, have been deposited in NCBI’s Gene Expression Omnibus and are accessible through GEO Series accession number GSE209557.

### Real-time quantitative PCR validation of targets

cDNA was synthesised from 175 ng of RNA using the Quantitect reverse transcription kit (205311, Qiagen). An aliquot of cDNA equivalent to 4.3 ng of the starting RNA template was used in duplicate for quantitative PCR. Quantitative PCR was conducted in 10 μl reaction volumes with specific primers for target genes and PowerUp SYBR Green master mix (A25742, Thermo Fisher Scientific) on an StepOnePlus Real-Time PCR system (Applied Biosystems). The internal control gene used for these analyses was the housekeeping gene Glyceraldehyde-3-phosphate dehydrogenase (*Gapdh*) and for relative quantification of gene expression the 2^−ΔΔCT^ method was used. Primers sequences included: *Avp* (5⍰-TGCCTGCTACTTCCAGAACTGC-3⍰ and 5⍰-AGGGGAGACACTGTCTCAGCTC-3⍰), caprin family member 2 (*Caprin2*, 5’-CAGGGTTAAGTGCAAGCGAT-3’ and 5’-CTGGTGGTTGACTGGTTGAG-3’), Cocaine And Amphetamine Regulated Transcript (*Cartpt*, 5’-CCTACTGCTGCTGCTACCTT-3’ and 5’-TCCACGGCAGAGTAGATGTC-3’), Cellular retinoic acid-binding protein 1 (*Crabp1*, 5’- TTTCCTCCACACACCTCTCC-3’ and 5’-CAGATGCCAAACCAGGGATG-3’), Galanin (*Gal*, 5’- CCTAGAGCAGTCCTGAGACC-3’ and 5’-TGAGGAGTTGGCGGAAGATT-3’), *Gapdh* (5’-ATGATTCTACCCACGGCAAG-3’ and 5’-CTGGAAGATGGTGATGGGTT-3’), glucagon-like peptide-1 receptor (*Glp1r*, 5’-GTTCCGCTGCTGTTCGTTAT-3’ and 5’- GCAAGCGTATGATGAGCCAA-3’), 5-Hydroxytryptamine Receptor 2A (*Htr2a*, 5’- GCCCGAGGCTCTACCATAAT-3’ and 5’- ACCCTTCACAGGAGAGGTTG-3’), *Opn3* (5’-CGACTGACAGGGACTCATCA-3’ and 5’- ATGGGACAGGCCAAAGAAGA-3’), *Oxt* (5’-TGCCCCAGTCTTGCTTGCT-3’ and 5’- TCCAGGTCTAGCGCAGCCC-3’), Prodynorphin (*Pdyn*, 5’-CCAGCCCCATCTCCTTAACT-3’ and 5’-AGACTGTTCCCCCTCGGTAT-3’), Pro-Melanin Concentrating Hormone (*Pmch*, 5’- TCGGTTGTTGCTCCTTCTCT-3’ and 5’-GATTCTGCTTGGAGCCTGTG), ras-related dexamethasone induced 1 (*Rasd1*, 5’-CCCTCAGCGTTGTGCCTACT-3’ and 5’-AAAGAGCGCACGGAACATCT-3’), and Transthyretin (*Ttr*, 5’- GACTCTGGTCATCGCCACTA-3’ and 5’-TTTGGCAAGATCCTGGTCCT-3’).

### Immunofluorescence

Two weeks following unilateral administration of AAV-shOpn3-eGFP in one SON and AAV-shNT-eGFP in the other SON, rats were deeply anesthetised with intraperitoneal administration of sodium pentobarbitone (100 mg/kg) and transcardially perfused with PBS followed by 4% (w/v) paraformaldehyde (PFA) in PBS. The brain was extracted, post-fixed in 4% (w/v) PFA overnight, cryoprotected in 30% (w/v) sucrose in PBS and frozen over liquid nitrogen. All sample collections were performed between 9.00 am and 12.00 pm. Coronal sections of the forebrain containing the hypothalamus were cut on a cryostat at 40 μm. Nonspecific protein binding was prevented by blocking the sections in PBS containing 0.3% (v/v) Triton X-100, 4% (w/v) bovine serum albumin (BSA), and 5% (v/v) donkey serum at RT for 1 hour. Following this, sections were incubated overnight at 4°C with primary antibodies against AVP neurophysin II (NP-II; MERCK, MABN845, PS41; 1:200); OXT NP-I (PS38; 1:100; Ben-Barak et al. (1985); CAPRIN2 (Proteintech, 20766-1-AP, 1:500), PDYN (Proenkephalin B (D-6); Santa Cruz Biotechnology, sc-398808, 1:100); CART (Bio-techne, AF163, 1:200); TTR (Bio-techne, NBP2-41101, 1:200), 5-HT2A (Alomone labs, ASR-033, 1:200). Sections were then washed in PBS and incubated with donkey anti-mouse Alexa Fluor 594 (Molecular Probes, A-21203) or donkey anti-rabbit Alexa Fluor Plus 647 (Molecular Probes, A32795) at a 1:500 dilution in PBS containing 0.1% (v/v) Triton, 4% (w/v) BSA, and 1% (v/v) donkey serum at RT for 1 h. Following this, the sections were washed in PBS, incubated with 4’,6-Diamidino-2-phenylindole dihydrochloride (DAPI, D1306; Molecular Probes) in PBS and mounted with ProLong Gold Antifade Mountant (Thermo Fisher Scientific, P36930).

### Image acquisition and data analysis

Images from RNAscope *in situ* hybridization studies were acquired with a Leica SP5-II AOBS confocal laser scanning microscope attached to a Leica DMI 6000 inverted epifluorescence microscope using a 63x PL APO CS lens with a 3.4-zoom factor. Quantification of *Opn3* RNA dots in the nucleus (DAPI labelling close to either AVP- or OXT-positive cytoplasm) or cytoplasm (AVP or OXT labelling) of AVP or OXT neurones was performed using a modular workflow plugin for Fiji created by Dr Stephen J Cross from the Wolfson Bioimaging Facility of the University of Bristol, as described (Barez-Lopez et al., 2022). Images from immunofluorescence studies were acquired with a Leica SP8 AOBS confocal laser scanning microscope attached to a Leica DM I8 inverted epifluorescence microscope using a 40x HC PL APO CS2 lens. Raw image files were processed to generate composite images using Fiji.

### Physiological analysis

The week following the delivery of bilateral administration of either AAV-shOpn3-eGFP or AAV-shNT-eGFP to the SON, food and water intake was measured daily for 5 weeks by weighing the amount of food and water available from each singly housed rat. Food and water intake was calculated by subtracting the obtained measure to the one from the previous day. Once a week, food and water intake were monitored every 4 hours during the dark phase of the diurnal cycle (4, 8 and 12 hours following the start of the dark phase) and following the 12 hours of the light phase of the diurnal cycle.

Following this, body temperature and motor activity were recorded by NanoTag (56747, Stoelting Europe) devices. To this aim, rats were anaesthetised with isoflurane (4% for induction, 2% for maintenance in O_2_) and nanotags were implanted subcutaneously in the lower edge of the scapula. Rats were allowed to recover from surgery for 3 days then nanotags started recording body temperature and motor activity at 5 min intervals for 3 days in basal conditions, 2 days of WD and 3 days of rehydration.

### Statistical analysis

Data are presented as mean ± SD. Statistical analyses were performed with GraphPad Prism software v8. Means between two groups were compared by 2-tailed unpaired Student’s t-test for RNAscope *in situ* hybridization quantification and for food and water intake measures every four hours. Means between two groups were compared by paired Student’s t-test for quantitative PCR outputs. Differences between means for more than two groups was performed by one-way analysis of variance (ANOVA) and the Tukey’s post hoc test to correct for multiple comparisons for RNAscope *in situ* hybridization quantifications. Analysis of physiological parameters was performed by two-way ANOVA and Sidak’s post hoc test. For motor activity data, the area under the curve (AUC) for the dark phase and for the light phase was calculated for each day, before assessing differences between AUC means by two-way ANOVA and Sidak’s post hoc test. Statistical significance was set at p⍰<⍰0.05.

## RESULTS

### Opn3 mRNA is expressed in SON AVP and OXT neurons and increases in response to stimulation

In order to get further information regarding the cell types expressing *Opn3* in the SON and its responses to stimuli, we performed RNAscope *in situ* hybridization labelling *Opn3* in combination with *Avp* and *Oxt* to identify AVP and OXT MCNs, respectively. We explored *Opn3* expression in the SON of euhydrated control rats, rats following 72-hours of WD, and 4, 8 and 24 hours of rehydration after WD.

Analysis of *Opn3* transcript expression in euhydrated conditions revealed that *Opn3* is found in both cytoplasmatic and nuclear compartments of AVP and OXT MCNs to the same extent (**Figure 1A, B**; t(6)=1.294, p = 0.2433, unpaired t-test in cytoplasm and t(6) = 0.3607, p = 0.7306, unpaired test in nucleus). WD increased *Opn3* abundance in the cytoplasm and nucleus of both AVP and OXT neurones (**Figure 1A, C, E**; AVP cytoplasm F (4, 15) = 4.113, p = 0.0191, one-way ANOVA, Tukey post-hoc test control vs WD p = 0.0116; AVP nucleus F (4, 15) = 6.815, p = 0.0025, one-way ANOVA, Tukey post-hoc test control vs WD p = 0.0030; OXT cytoplasm F (4, 15) = 3.633, p = 0.0292, Tukey post-hoc test control vs WD p = 0.0203; OXT nucleus F (4, 15) = 4.850, p = 0.0104, Tukey post-hoc test control vs WD p = 0.0072). There were no differences in the expression of *Opn3* between AVP and OXT neurons following WD (**Figure 1A, D**; t(6) = 0.5622, p = 0.5944, unpaired t-test in cytoplasm and t(6) = 0.3607, p = 0.7306, unpaired t-test in nucleus). During rehydration, *Opn3* expression decreased towards control levels in the cytoplasm and nucleus of AVP and OXT cells as early as 4 hours following water reintroduction (**Figure 1A, C, E**).

**Figure 1.**
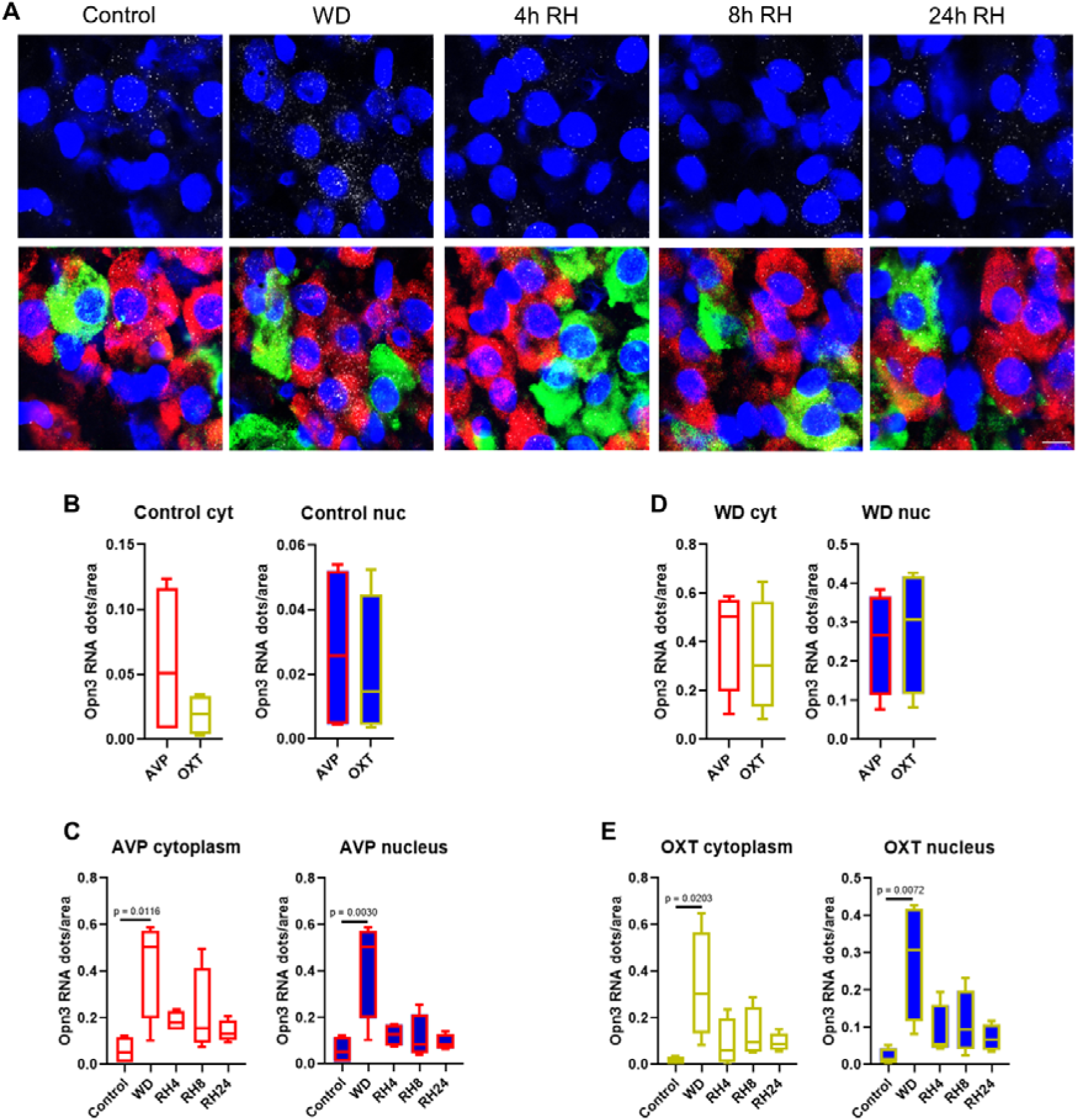
*Opn3* mRNA expression in the supraoptic nucleus (SON). DAPI (blue), *Avp* (red), *Oxt* (green) and *Opn3* (grey) labelled by RNAscope *in situ* hybridisation in control (n = 4) conditions, after 72 hours of water deprivation (WD; n = 4), after 72 hours of WD followed by 4 hours rehydration (4h RH; n = 4), 8 hours rehydration (8h RH; n = 4) or 24 hours rehydration (24 h RH; n = 4). (**A**). Graphs showing gene expression as a function of *Opn3* mRNA dots/cytoplasm of AVP vs OXT neurones in control conditions (**B**), *Opn3* mRNA dots/nuclei of AVP vs OXT neurones in control conditions (**C**), *Opn3* mRNA dots/cytoplasm of AVP vs OXT neurones in WD conditions (**D**), *Opn3* mRNA dots/nuclei of AVP vs OXT neurones in WD conditions (**E**), *Opn3* mRNA dots/cytoplasm AVP neurones (**F**), *Opn3* mRNA dots/nuclei AVP neurones (**G**), *Opn3* mRNA dots/cytoplasm OXT neurones (**H**), *Opn3* mRNA dots/nuclei OXT neurones (**I**). Data are expressed as mean ± SD. Scale bar represents 10 µm.

### Opn3 regulates peptide production in the SON

As a first step to unravel the functions of Opn3 in the SON, we performed transcriptomic analysis in *Opn3* gene knockdown rat SONs (Opn3KD). We knocked down the expression of *Opn3 in vivo* by delivering AAV vectors containing a specific *Opn3* shRNA and an eGFP tag unilaterally into one SON by stereotaxic brain surgery. As controls, AAVs delivering a non-target scrambled shRNA and an eGFP tag were delivered unilaterally to the opposite SON in the same animals (**Figure 2A**). The delivery of Opn3KD vectors significantly decreased *Opn3* mRNA expression between 55 and 89% in comparison to control SONs (**Figure 2B**). We then sequenced the polyadenylated transcriptomes of control and Opn3KD SONs by RNA-seq. Principal component analyses (PCA) using all genes revealed a distinct separation between the transcriptomes of control and Opn3KD SONs. Principal component 1 (PC1) explained 73% of total variance that was attributable to the experimental condition (**Figure 2C**). Differential expression analysis identified a total of 3711 differentially expressed genes (DEGs) (P_Adj_ < 0.05), 1986 of which were upregulated whilst 1725 were downregulated (**Figure 2D, Table 2-1**).

**Figure 2.**
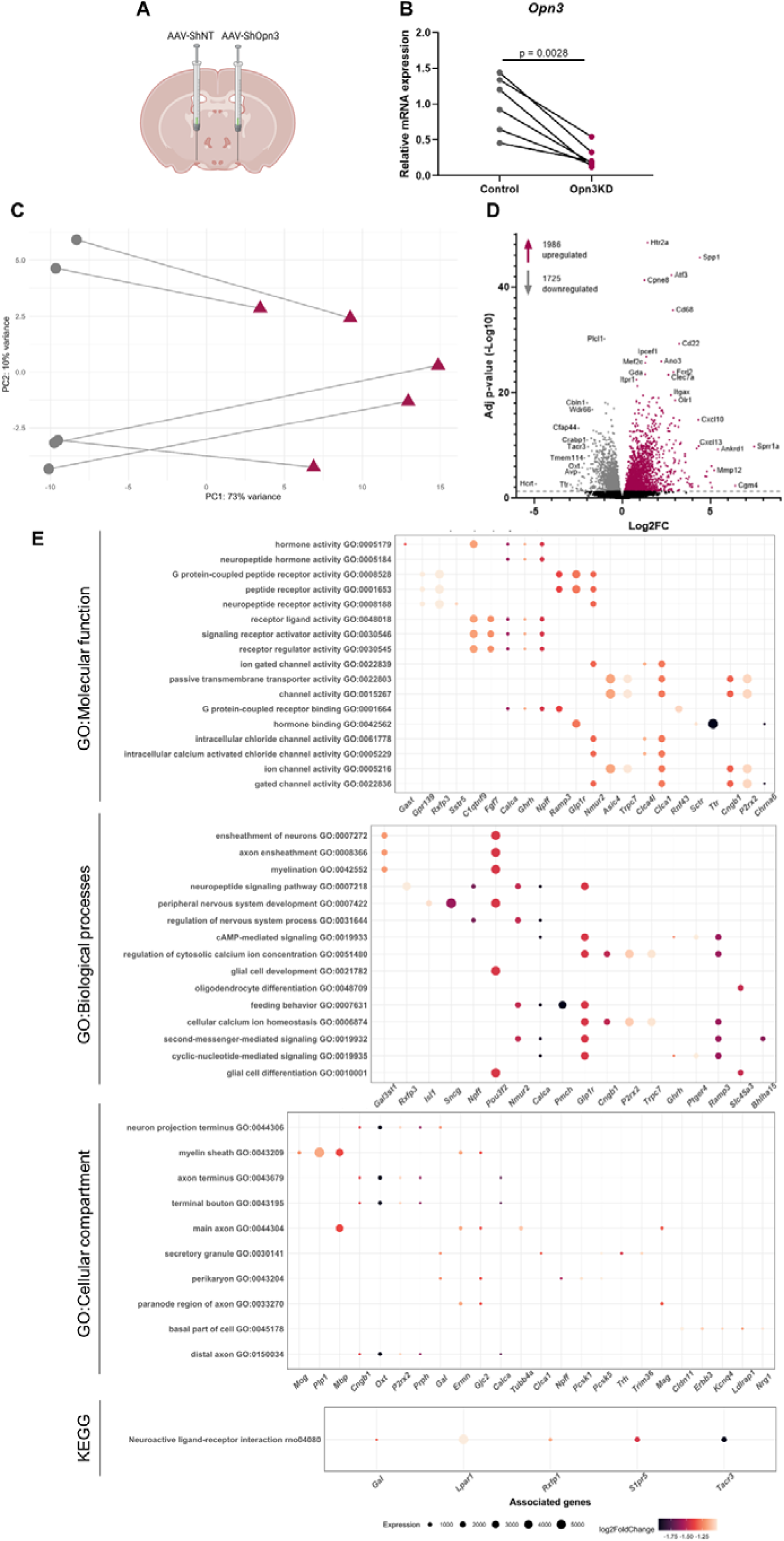
Gene profiling analysis by RNA-seq in Opn3 knockdown supraoptic nucleus (SON). Opn3 was knocked down by delivering AAV vectors containing a specific *Opn3* shRNA (Opn3KD, n = 5) unilaterally into one SON and AAVs delivering a non-target scrambled shRNA to the opposite SON as a control (n = 5; **A**). *Opn3* mRNA expression by quantitative PCR in control and Opn3KD SONs showing *Opn3* knock down efficiency (**B**). Principal component analysis showing distinct separation between control and Opn3KD conditions (**C**). Volcano plot of Opn3KD vs control SON transcriptomes showing 1986 upregulated (purple) and 1725 downregulated (grey) genes (P_Adj_ < 0.05, **D**). Pathway analysis of changes in the SON transcriptome resulting from *Opn3* knock down using GO and KEGG databases. Dot plot of all enriched terms retrieved for each category ranked according to PAdj value from top to bottom in increasing order. The topmost significant associated differentially expressed proteins of each over-represented category are shown as dots coloured based on Log2FC and sized according to transcript abundance measured by normalised read counts aligned to each gene (**E)**.

In order to obtain insights regarding the physiological role of Opn3 in the SON, we performed pathway analysis of all the DEGs between control and Opn3KD samples by interrogating GO and KEGG databases. This analysis retrieved broad terms such as “protein binding” or “ion binding” in GO:Molecular functions category and “positive regulation of biological processes” or “localisation” in the GO:Biological processes category (*Figure 2-1, Table 2-2*). To further explore the consequences from knocking down *Opn3*, we performed pathway analysis on those downregulated DEGs (P_Adj_ < 0.005) with a baseMean > 10 and log2 fold-change (log2FC) < -1. This analysis retrieved 19 enriched terms in the GO:Molecular function (GO:MF) category with “hormone activity” and “neuropeptide hormone activity” being the most significant (*Figure 2E, table 2-3*). The GO:Biological processes

**Figure 2-1.**
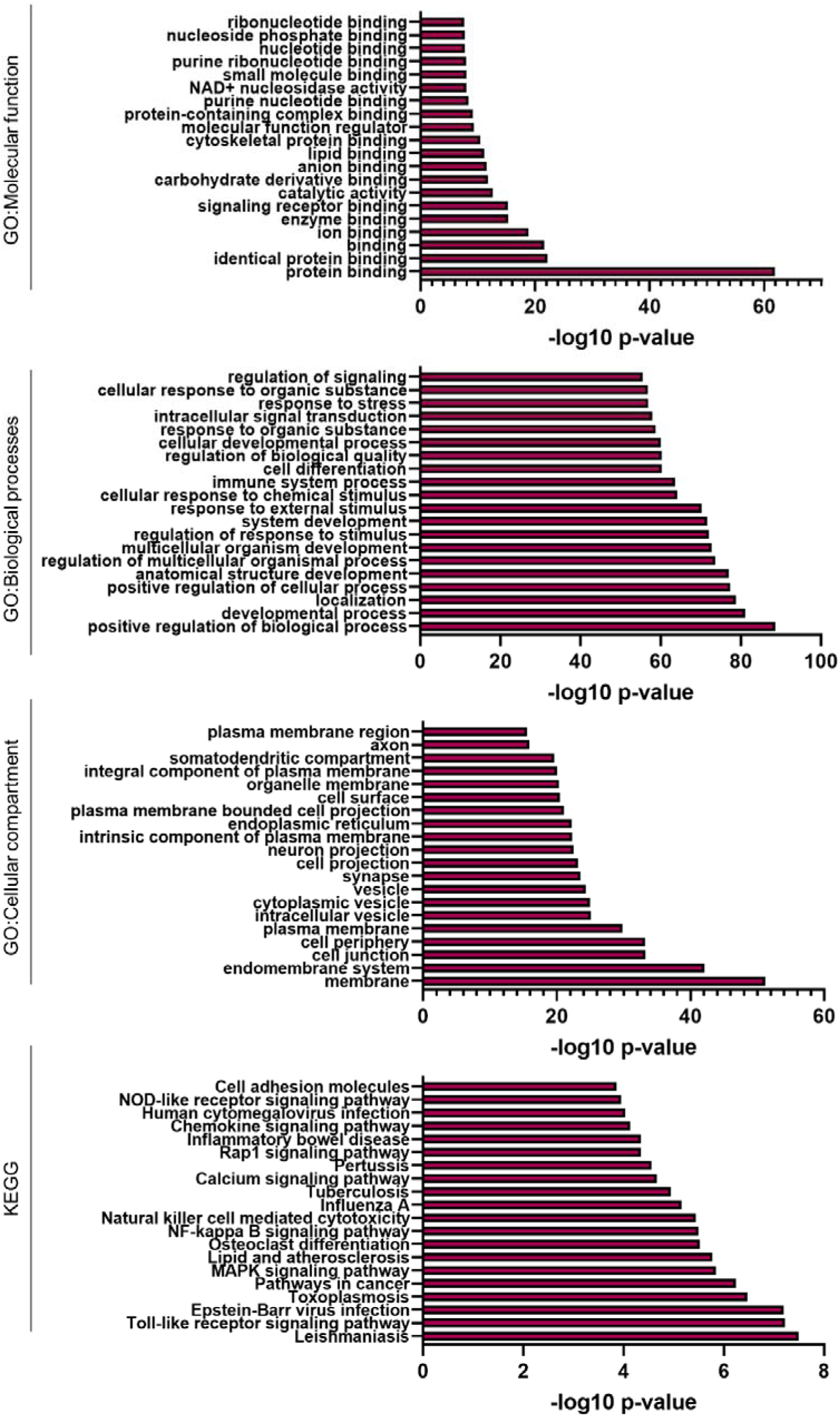
Pathway analysis of all the DEGs with base mean > 10 between control and Opn3KD samples using GO and KEGG databases.

(GO:BP) category retrieved 65 significantly enriched terms including “ensheathment of neurons”, “neuropeptide signalling pathway” or “feeding behaviour” (**Figure 2E, table 2-3**). The GO:Cellular compartment (GO:CC) category highlighted 10 enriched terms such as “neuron projection terminus” and “secretory granule” (**Figure 2E, table 2-3**). Kyoto Encyclopaedia of Genes and Genomes (KEGG) analysis retrieved the single significant term “Neuroactive ligand-receptor interaction” (**Figure 2E, table 2-3**).

All pathway analysis categories were indicative of Opn3 mediating a role in neuropeptide-related pathways. In view of this, we validated by qPCR a subset of genes included in the “neuropeptide hormone activity”, “neuropeptide signalling pathway”, “secretory granule” and “neuroactive ligand-receptor interaction” terms from the GO:MF, GO:BP, GO:CC and KEGG categories, respectively. These genes included *Avp, Cartpt, Gal, Glp1r, Oxt, Pdyn*, and *Pmch*. Due to their pivotal role in osmoregulation (Konopacka et al., 2015; Greenwood et al., 2016), we also included the genes *Caprin2* and *Rasd1*. Since Opn3 can bind retinol to mediate light-sensitive actions, we also studied the expression of *Ttr* and *Crabp1* which encode proteins that bind and transport retinol and the retinol derivative retinoic acid, respectively. All these genes were significantly downregulated (P_Adj_ < 0.005) in Opn3KD samples (**table 2-1**). In addition, we also studied the expression of *Hrt2a*, encoding the serotonin receptor 5-HT2A, due to its important physiological functions and because it was the most significant upregulated gene in Opn3KD samples (**Figure 2D, table 2-1**). The changes in the expression of these genes validated the RNA-seq output, with the exception of *Pmch*, which downregulation in Opn3KD SONs was not statistically significant (**Figure 3**; t(4) = 4.795, p = 0.0087, paired t-test *Avp*; t(4) = 4.748, p = 0.0090, paired t-test *Caprin2*; t(4) = 6.011, p = 0.0039, paired t-test *Cartpt*; t(4) = 4.756, p = 0.0089, paired t-test *Crabp1*; t(4) = 6.902, p = 0.0023, paired t-test *Gal*; t(3) = 3.422, p = 0.0418, paired t-test *Glp1r*; t(4) = 7.066, p = 0.0021, paired t-test *Htr2a* ; t(4) = 8.546, p = 0.0010, paired t-test *Oxt*; t(4) = 4.532, p = 0.0106, paired t-test *Pdyn*; t(4)=2.095, p = 0.1042, paired t-test *Pmch*; t(4) = 3.510, p = 0.0247, paired t-test *Rasd1*; t(4) = 2.825, p = 0.0476, paired t-test *Ttr*). This was probably due to the high variability in the number of *Pmch* reads in control samples, which had a base mean of 423.80 and a standard deviation of 444.05.

**Figure 3.**
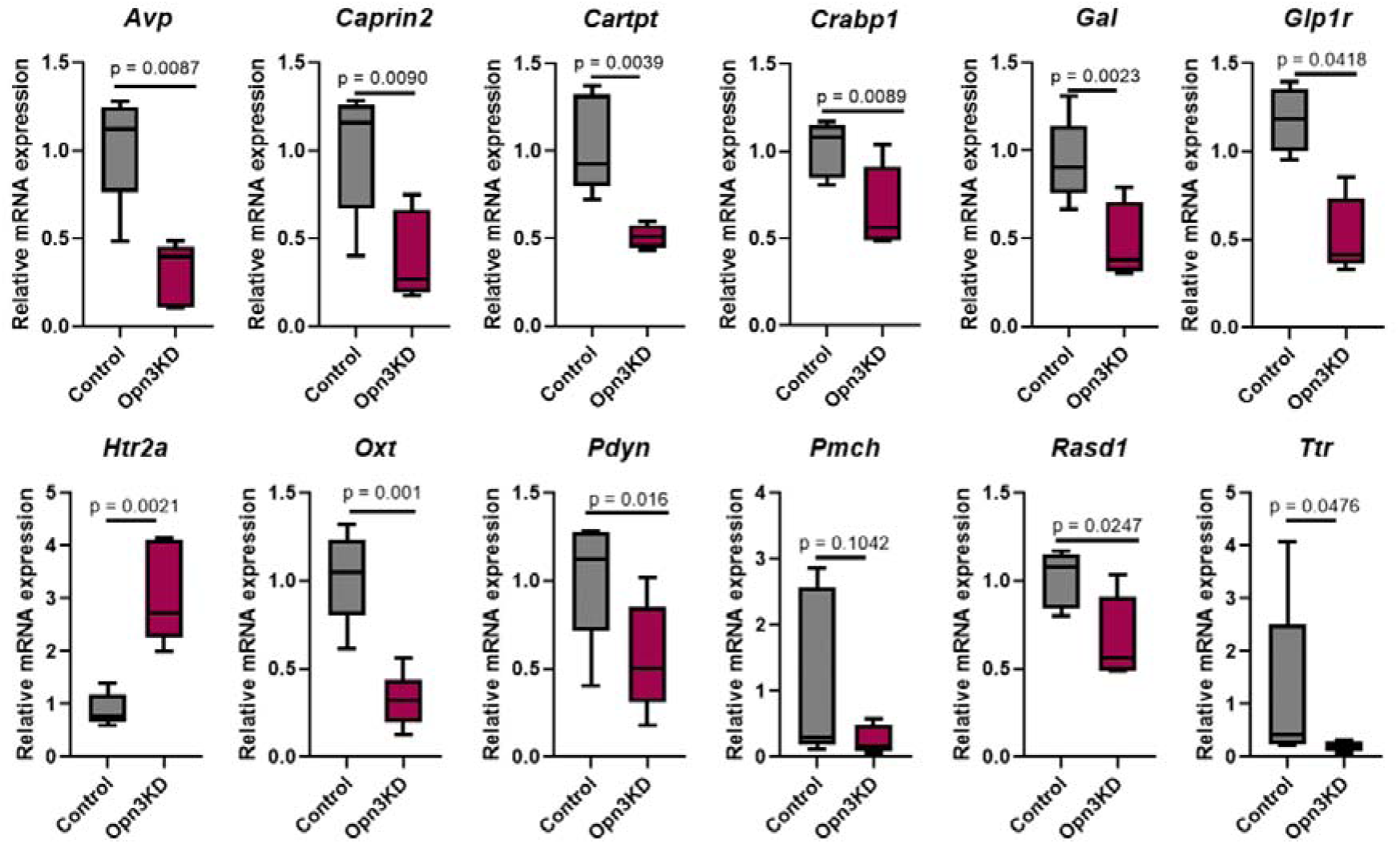
Quantitative PCR validation of mRNA expression in the supraoptic nucleus (SON) of control (n = 5) and Opn3KD (n = 5) rats. Arginine vasopressin (*Avp*), Caprin family member 2 (*Caprin2)*, Cocaine and Amphetamine Regulated Transcript (*Cartpt)*, Cellular retinoic acid-binding protein 1 (*Crabp1*), Galanin (*Gal*), glucagon-like peptide-1 receptor (*Glp1r*), 5-Hydroxytryptamine Receptor 2A (*Htr2a*), oxytocin (*Oxt*), Prodynorphin (*Pdyn*), Pro-Melanin Concentrating Hormone (*Pmch*), ras-related dexamethasone induced 1 (*Rasd1*), and Transthyretin (*Ttr*) were corrected for Glyceraldehyde-3-phosphate dehydrogenase (*Gapdh*) RNA content. Data are expressed as mean ± SD.

To assess the effects of the downregulation of these transcripts on their encoded proteins, we performed immunofluorescence analysis on brain samples of rats that had been injected unilaterally with AAVs delivering Opn3 shRNA (Opn3KD) and an eGFP tag in one SON and with non-target scrambled shRNA and an eGFP tag (control) into the other SON. Successful AAV delivery was confirmed by GFP expression in the SON (**Figure 4A, A’**). There was a remarkable decrease in the staining for the neuropeptides AVP and OXT in Opn3KD samples in comparison to their corresponding controls, indicating a decrease in protein production (**Figure 4B, B’, C, C’**). The staining of Caprin family member 2 (Caprin2, encoded by *Caprin2*), an RNA binding protein which binds *Avp* and *Oxt* RNAs increasing their transcript stability (Konopacka et al., 2015; Barez-Lopez et al., 2022), was also reduced (*Figure 4D, D’*). Staining of neuropeptide Proenkephalin-B (PDYN, encoded by *Pdyn*), did not differ between control and Opn3KD samples suggesting that, at least in basal conditions, PDYN production is not altered in the SON (**Figure 4E, E’**). Similarly, staining of the neuropeptide Cocaine- and amphetamine-regulated transcript protein (CART, encoded by *Cartpt*), was detected in some neurones and neuronal processes in the SON, but did not differ between control and Opn3KD samples (**Figure AF, F’**). Transthyretin (TTR, encoded by *Ttr*), displayed less staining in Opn3KD SON in comparison to control SON, indicating decreased TTR production (**Figure 4G, G’**). The 5-hydroxytryptamine receptor 2A (5-HT2A, encoded by *Htr2a*) displayed no obvious differences in the staining intensity between control and Opn3KD SONs, even though the transcript levels were highly increased (**Figure 4H, H’**).

**Figure 4.**
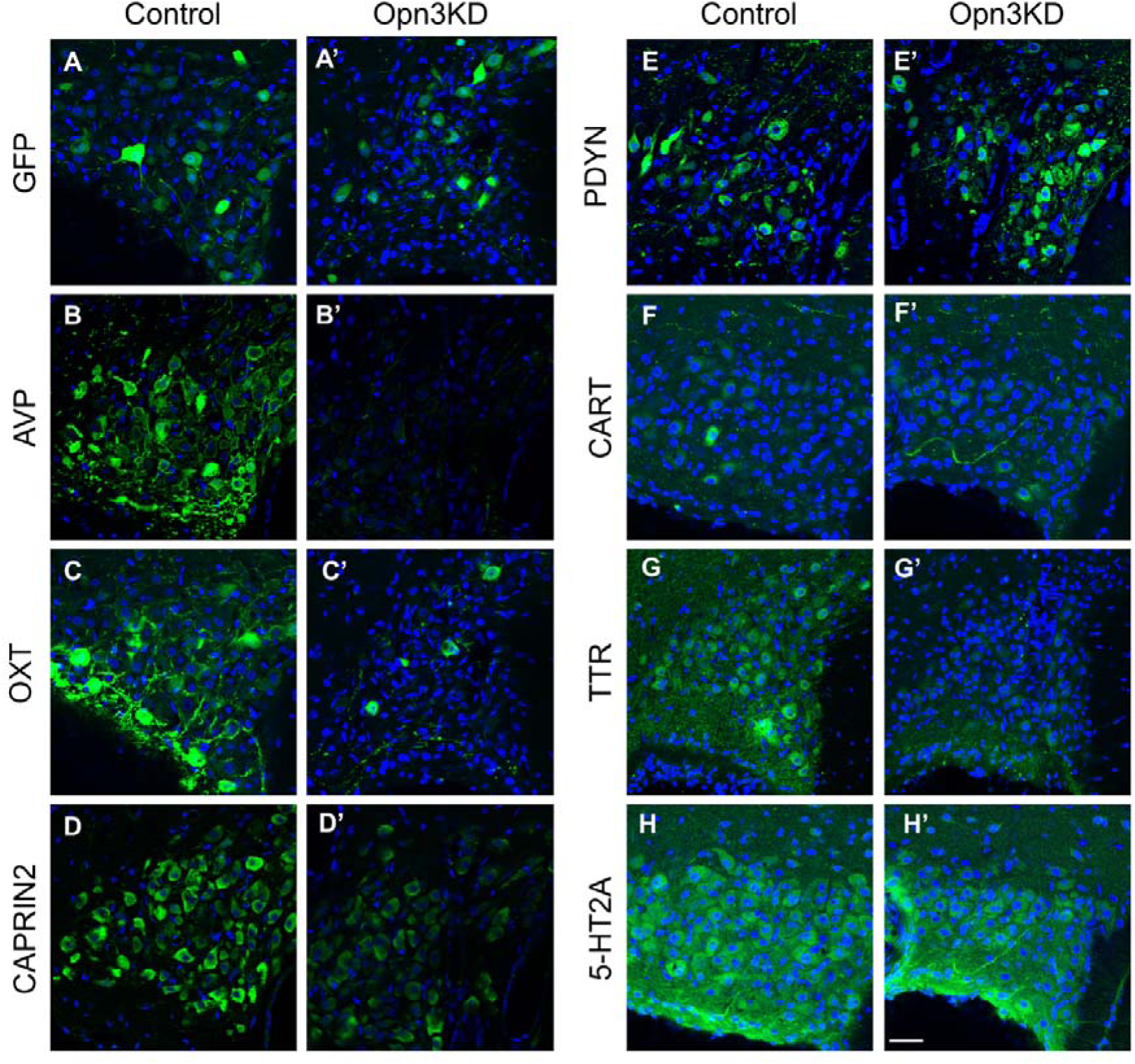
Immunofluorescence of supraoptic nucleus (SON) of control and Opn3KD rats. Detection of green fluorescent protein (GFP), arginine vasopressin (AVP), oxytocin (OXT), Caprin family member 2 (CAPRIN2), Prodynorphin (PDYN), Cocaine- and amphetamine-regulated transcript protein (CART), Transthyretin (TTR) and Serotonin 5-HT2A receptor (5-HT2A). Images are representative of n = 4. Scale bar represents 40 µm.

### Knocking down Opn3 in the SON alters water intake, body temperature and motor activity

Since knocking down *Opn3* in the SON led to the downregulation of genes encoding peptides that mediate water homeostasis, food intake, locomotion, and sleep among other processes, we studied the physiological effects of knocking down *Opn3*. We focused on physiological effects related to food and water intake, body temperature, and motor activity. To this end, we knocked down Opn3 in the SON of rats by delivering AAV-shOpn3-eGFP bilaterally into both SONs by stereotaxic brain surgery (Opn3KD rats). Rats injected bilaterally with AAV-shNT-eGFP into both SONs were used as controls (**Figure 5A**). Successful Opn3 downregulation was assessed at termination by analysing *Opn3* expression by qPCR that revealed an average of 78% KD efficiency (t(24) = 7.212, p <0.0001, unpaired t-test; **Figure 5B**). During the week following the delivery of the AAVs, water and food intake was measured daily for 5 weeks and the average weekly water and food intake per 100 g of body weight (b.w.) is presented. We found a significant difference in water intake between weeks (F (4, 55) = 3.203, p = 0.0196, two-way ANOVA) and between control and Opn3KD rats (F (1, 55) = 16.74, p = 0.0001, two-way ANOVA), though the interaction between these terms was not significant (F (4, 55) = 2.079, p = 0.0960, two-way ANOVA; **Figure 5C**). Sidak’s multiple comparisons test also revealed significant pairwise differences between control and Opn3KD rats in water intake in week 3 (p = 0.0435) and week 4 (p = 0.0478). Food intake was different between weeks (F (4, 55) = 51.41, p < 0.0001, two-way ANOVA) but there were no differences between control and Opn3KD rats (F (1, 55) = 0.6874, p = 0.4106, two-way ANOVA) and neither the interaction between these terms was significant (F (4, 55) = 0.2178, p = 0.9274, two-way ANOVA; **Figure 5D**). In order to get a better understanding regarding water and food intake, once a week food and water intake were assessed every 4 hours during the dark phase and after the entire 12-h light phase. As already reported (Zucker, 1971), most of the water and food intake took place during the dark phase (**Figure 5E**). Water intake was significantly higher in Opn3KD rats during the light phase in week 2 (t(11) = 2.706, p = 0.0204, unpaired t-test), in week 3 from 4 to 8 hours into the dark phase (t(11) = 2.459, p = 0.0318, unpaired t-test) and during the light phase (t(11) = 2.226, p = 0.0478, unpaired t-test), and in week 5 from 4 to 8 hours into the dark phase (t(11) = 2.219, p = 0.0484, unpaired t-test) and 8 to 12 hours into the dark phase (t(11) = 2.496, p = 0.0297, unpaired t-test; **Figure 5E**), confirming that water intake is higher in Opn3KD rats. Food intake was not significantly different between control and Opn3KD rats during the different phases of the diurnal cycle in any of the weeks (*Figure 5E*).

**Figure 5.**
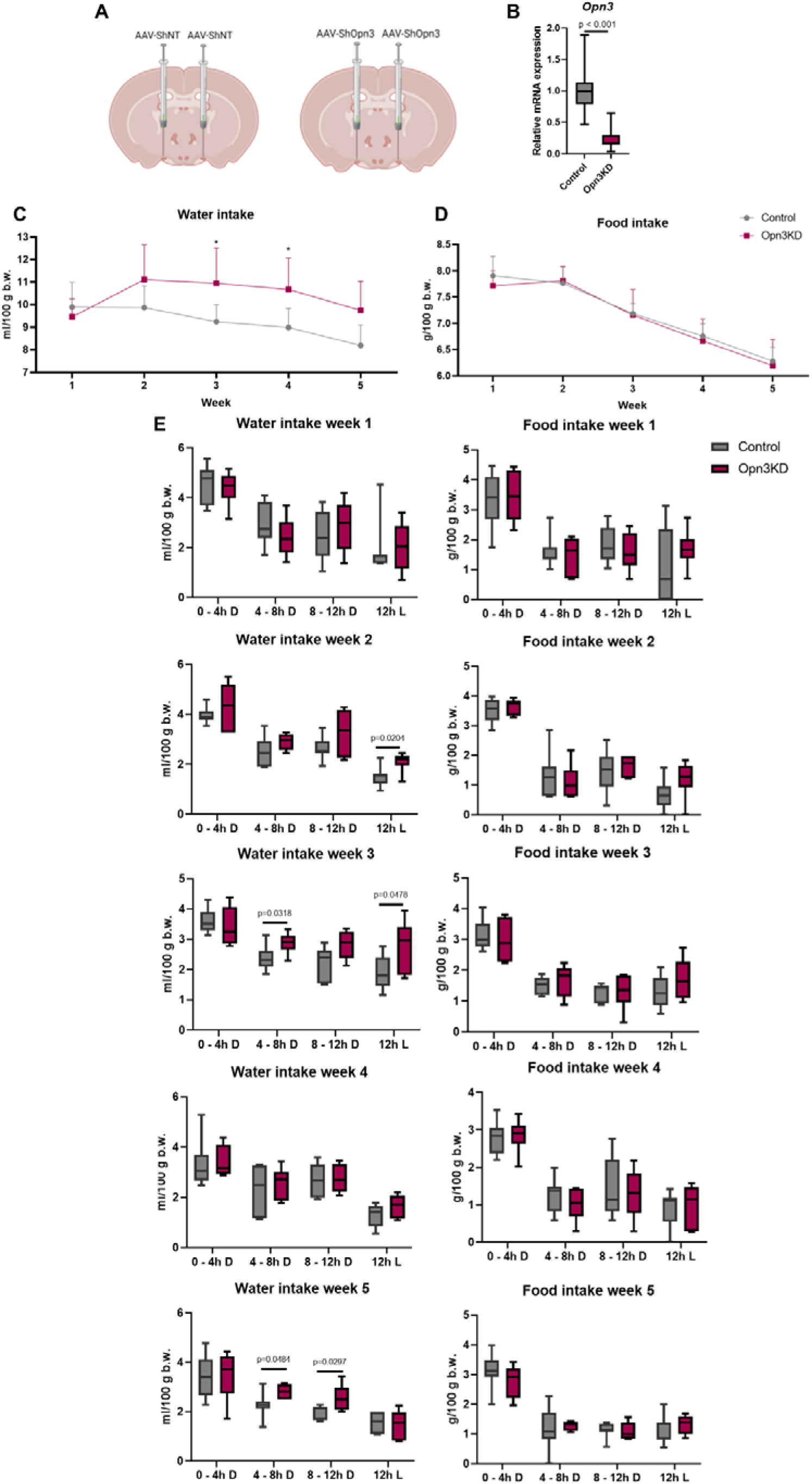
Food and water intake in control and Opn3KD rats. Opn3 was knocked down by delivering AAV vectors containing a specific *Opn3* shRNA (Opn3KD, n = 6) bilaterally into both SONs. AAVs delivering a non-target scrambled shRNA were administered bilaterally to both SONs as controls (n = 5; **A**). *Opn3* mRNA expression by quantitative PCR in control and Opn3KD SONs showing *Opn3* knock down efficiency (**B**). Water (**C**) and food (**D**) intake were measured daily for 5 weeks. Once a week water and food intake was measured every 4 hours during the dark (D) phase of the diurnal cycle and following the 12 hours of the light (L) phase of the diurnal cycle (**E**). Data are expressed as mean ± SD.

Following water and food intake measurements, body temperature and motor activity were recorded through the insertion of a nanotag activity measuring device. Body temperature and motor activity were recorded every five minutes for 3 consecutive days in basal conditions (B1, B2 and B3) and, since the data was indicating that OPN3 is involved in water homeostasis, during 2 days of WD (WD1, WD2) followed by 3 days of rehydration (RH1, RH2 and RH3). Body temperature recordings and motor activity measurements were averaged for the dark phase and the light phase each day. Knocking down *Opn3* in the SON did not impair the circadian rhythmicity observed in body temperature and motor activity between the dark and the light phase (**Figure 6A, B**). In the dark phase, body temperature was not significantly different between days (F(7, 88) = 0.8426, p = 0.5551, two-way ANOVA) or between control and Opn3KD animals (F (1, 88) = 3.559, p = 0.0625, two-way ANOVA). However, the interaction between these two terms was significant (F (7, 88) = 4.396, p = 0.0003, two-way ANOVA; **Figure 6C**), indicating that the response of body temperature to osmotic stimulation is different between control and Opn3KD rats. In addition, a Sidak post-hoc test revealed significant pairwise differences between control and Opn3KD rats in body temperature on day B1 (p = 0.0043) and B3 (p = 0.0321), where Opn3KD rats had higher body temperature values (**Figure 6C**). In the light phase, body temperature was significantly different between days (F (7, 88) = 2.673, p = 0.0148, two-way ANOVA) but it did not differ between control and Opn3KD animals (F (1, 88) = 1.117, p = 0.2935, two-way ANOVA; **Figure 6D**). The interaction between body temperature in days and in control or Opn3KD animals was not significant (F (7, 88) = 0.09697, p = 0.9984, two-way ANOVA; **Figure 6D**).

**Figure 6.**
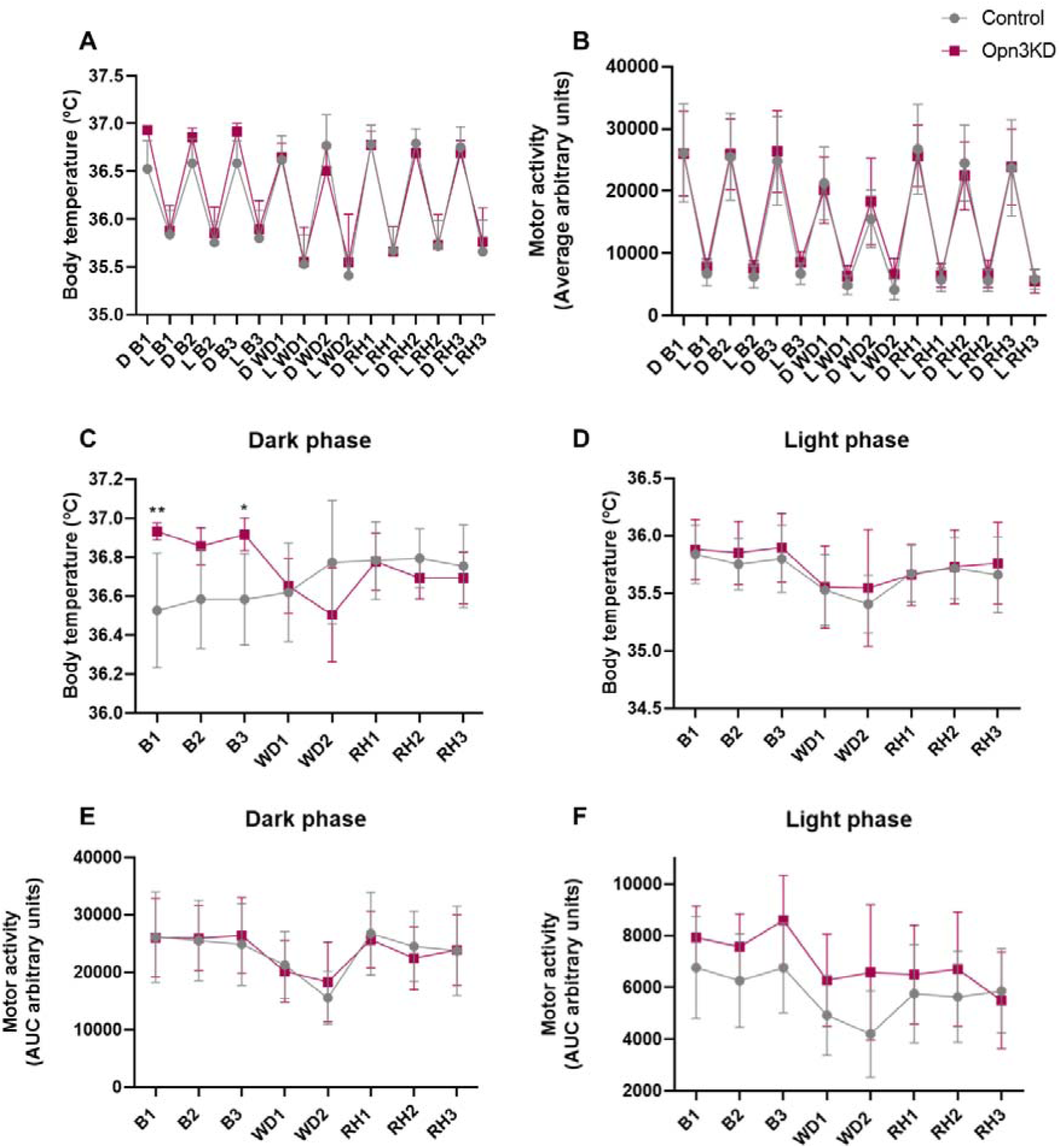
Body temperature and motor activity in control and Opn3KD rats. Body temperature (**A**) and motor activity (**B**) oscillations between the dark (D) and the light (L) phase of the diurnal cycle in basal conditions (B1, B2, B3), water deprivation (WD1, WD2) and rehydration (RH1, RH2, RH3). Body temperature in the dark (**C**) and the light phase (**D**). Motor activity in the dark (**E)** and the light phase (**F**). Data are expressed as mean ± SD.

Motor activity measurements in the dark phase revealed significant differences between days (F (7, 88) = 3.309, p = 0.0036, two-way ANOVA), but there were no differences between control and Opn3KD rats (F (1, 88) = 0.003026, p = 0.9563, two-way ANOVA) and the interaction between these two terms was not significant (F (7, 88) = 0.1870, p = 0.9875, two-way ANOVA; **Figure 6E**). In the light phase, there were significant differences in motor activity between days (F (7, 88) = 2.857; p =0.0098, two-way ANOVA) and also between Opn3KD and control animals (F (1, 88) = 11.08; p = 0.0013), where Opn3KD animals were more active than controls during the light phase. The interaction between values on different days and in different animal groups was not significant (F (7, 88) = 0.6340, p = 0.7266; **Figure 6F**).

## DISCUSSION

In this work we have addressed, for the first time, the role of Opn3 in the mammalian brain, in the hypothalamic SON. *Opn3* mRNA expression and Opn3 protein localisation have been identified in the hypothalamic SON, and we have observed that *Opn3* expression is response to osmotic stimulation in this structure. We thus hypothesised that Opn3 mediates important functions in the SON.

As a first step to unravel the role of Opn3 in the SON, we identified the neuronal types expressing *Opn3*. We found that *Opn3* is present to the same extent in both AVP and OXT neurones and that its expression increases in both neuronal types to the same degree in response to osmotic stimulation. This suggests that Opn3 exerts a role in both AVP and OXT neurones in the SON. To obtain further insights regarding the role of Opn3 in the SON, we performed gene profiling experiments using RNA-seq by comparing control and Opn3KD rats. Pathway analysis revealed enriched terms related to neuropeptide signalling pathways. We found that several genes known to be involved in water homeostasis were amongst the DEGs identified in Opn3KD SON. These included the *Avp* and *Oxt* neuropeptide-encoding RNAs, which encode peptide hormones with well-known roles in the control of water reabsorption and natriuresis respectively at the level of the kidney (Breyer and Ando, 1994). In addition, the *Caprin2* mRNA is under Opn3 control. The Caprin2 protein regulates *Avp* and *Oxt* transcript polyadenylation and stability/translation (Konopacka et al., 2015; Barez-Lopez et al., 2022). Expression of the mRNA encoding small G protein Rasd1, that controls the transcriptional response to osmotic stress (Greenwood et al., 2016), is also downregulated by Opn3 knockdown. Whilst these and other genes that increase their expression in response to WD were downregulated in Opn3KD SONs, other genes that increase in expression in response to WD, such as *Azin1, Creb3l1, Giot1* or *Nr4a1* (Pauza et al., 2021), were not altered in Opn3KD animals. These findings, in addition to the increase in *Opn3* expression in the SON in response to hyperosmotic stimuli (Greenwood et al., 2015; Pauza et al., 2021), indicate that whilst Opn3 is involved in coordinating some aspects of the response to WD, other parallel processes are also engaged.

Other DEGs of interest that were downregulated by Opn3KD included the neuropeptides *Pdyn*, which is involved in motivation, response to stress, and pain sensitivity (Karkhanis & Al-Hasani, 2020); *Cartpt*, that mediates food intake regulation, neuroprotection, reward and cocaine-induced locomotion (Zhang et al., 2012; Ong and McNally, 2020); and *Gal*, which has been implicated in learning and memory, nociception, feeding, and sleep (Vrontakis, 2002; Kroeger et al., 2018), among many other functions. *Ttr*, encoding a transport protein that transports the thyroid hormone thyroxine and retinol (Liz et al, 2020), and *Crabp1*, encoding a protein that binds the retinol derivative retinoic acid (Napoli, 2016), were also downregulated in Opn3KD. Whether the lack of proteins binding retinol and its derivatives is related to the absence of Opn3, which would require retinol for photoactivation, is a matter that requires further investigation. Expression of *Hrt2a* was highly upregulated in Opn3KD, although the serotonin (5-HT) receptor 5-HT2A encoded protein did not appear to change its content in Opn3KD, at least in basal conditions. The increase of *Hrt2a* expression is of interest as 5-HT2A-mediated effects of 5-HT or 5-HT antagonists are involved in memory and cognition (Zhang and Stackman, 2015), psychiatric disorders (de Angelis, 2002), body temperature (Voronova et al., 2016) and sleep (Vanover and Davis, 2010). In addition, 5-HT is necessary for the increases in *Avp* and *Oxt* mRNA accumulation following hyperosmotic stimulation (Carter and Murphy, 1989), suggesting that increases in *Hrt2a* in Opn3KD SONs maybe a compensatory mechanism as a result of downregulation of *Avp* and *Oxt* mRNAs.

The findings above suggested that OPN3 is involved in the production of AVP, OXT and other neuropeptides, so downregulation of *Opn3* could be affecting behavioural and physiological parameters, including water and food intake, body temperature and motor activity. Water intake was increased in Opn3KD rats, while food intake was not different from controls. Increased water ingestion could possibly be related with alterations in water balance regulation. Even though Opn3KD rats ingested more fluid than controls, the general water and food intake patterns did not differ. Body temperature was also increased in Opn3KD animals, and the response of this parameter to WD was different between control and Opn3KD rats, but only in the dark phase of the diurnal cycle. Endogenous AVP in the brain has been suggested to mediate antipyretic effects (Richmond, 2003) and central infusion of AVP has been shown to produce hypothermia (Pittman et al., 1998), suggesting that increased body temperature in Opn3KD rats could be a consequence of decreased AVP production. Motor activity recordings were consistent with previous findings in control rats (Martelli et al., 2012), showing decreases in motor activity in response to WD in the dark phase. Interestingly, motor activity was increased in Opn3KD rats during the light phase, which is the inactive phase for nocturnal animals, and when rats are mostly asleep (Martelli et al., 2012), indicating that downregulation of *Opn3* increases motor activity during the diurnal nadir period and possibly disrupts sleep.

We have produced evidence that Opn3 in the SON is involved in the regulation of a number of neuropeptides and other proteins that participate in water homeostasis, body temperature, motor activity, and possibly sleep. It has long been known that AVP and OXT show daily rhythms of secretion (Forsling, 2000) that can be affected by constant light (Juszczak et al., 1995). The fact that Opn3 is a light-sensitive protein raises interesting possibilities, such as the presence of intrinsic circadian control mechanisms directly in the SON. Because body temperature and motor activity were altered in Opn3KD rats at specific phases of the diurnal cycle, it is possible that the mechanisms regulating these processes might be indeed light dependent. Further studies should aim to fully address if Opn3 in the mammalian brain plays a light-dependent or -independent function to get further insights into the physiological and pathological effects of light in the brain. In addition, the next step would be to address the potentially differential role of Opn in AVP and OXT neurones and to further explore the role of Opn3 in hyperosmotic stimuli and other SON-sensitive stimuli, such as lactation for example. To conclude, we provide the first insights into the role of Opn3 in the brain and identify the SON as an important target site for its action.

## ACKNOWLEGMENTS

We gratefully acknowledge the Wolfson Bioimaging Facility, in particular Dr Stephen J Cross, for their support and assistance in this work. This research was supported by grants from the Biotechnology and Biological Sciences Research Council (BBSRC; BB/R016879/1) to D.M., S.B.L., P.B. and M.P.G., from the Leverhulme Trust (RPG-2017-287) to D.M. and M.P.G., and a British Society for Neuroendocrinology Project Support Grant to S.B.L.

